# The Functional Significance of Bacterial Predators

**DOI:** 10.1101/2021.02.22.432408

**Authors:** Bruce A. Hungate, Jane C. Marks, Mary E. Power, Egbert Schwartz, Kees Jan van Groenigen, Steven J. Blazewicz, Peter Chuckran, Paul Dijkstra, Brianna K. Finley, Mary K. Firestone, Megan Foley, Alex Greenlon, Michaela Hayer, Kirsten S. Hofmockel, Benjamin J. Koch, Michelle C. Mack, Rebecca L Mau, Samantha N. Miller, Ember M. Morrissey, Jeff R. Propster, Alicia M. Purcell, Ella Sieradzki, Evan P. Starr, Bram W. G. Stone, César Terrer, Jennifer Pett-Ridge

## Abstract

Predation structures food webs, influences energy flow, and alters rates and pathways of nutrient cycling through ecosystems, effects that are well documented for macroscopic predators. In the microbial world, predatory bacteria are common, yet little is known about their rates of growth and roles in energy flows through microbial food webs, in part because these are difficult to quantify. Here, we show that growth and carbon uptake were higher in predatory bacteria compared to non-predatory bacteria, a finding across 15 sites, synthesizing 82 experiments and over 100,000 taxon-specific measurements of element flow into newly synthesized bacterial DNA. Obligate predatory bacteria grew 36% faster and assimilated carbon at rates 211% higher than non-predatory bacteria. These differences were less pronounced for facultative predators (6% higher growth rates, 17% higher carbon assimilation rates), though high growth and carbon assimilation rates were observed for some facultative predators, such as members of the genera *Lysobacter* and *Cytophaga*, both capable of gliding motility and wolfpack hunting behavior. Added carbon substrates disproportionately stimulated growth of obligate predators, with responses 63% higher than non-predators for the Bdellovibrionales and 81% higher for the Vampirovibrionales, whereas responses of facultative predators to substrate addition were no different from non-predators. This finding supports ecological theory that higher productivity increases predator control of lower trophic levels. These findings also indicate that the functional significance of bacterial predators increases with energy flow, and that predatory bacteria influence element flow through microbial food webs.

## Introduction

Bacteria that prey on other bacteria are too small to engulf their victims, yet they consume them no less ferociously. Members of the Bdellovibrionales attach to prey cells, penetrate the cell membrane, and then take up residence in the host cytoplasm, consuming cellular constituents while growing filaments and producing daughter cells that eventually lyse and kill the prey (*1*). Some bacterial predators have names that tell their mode of predation: *Vampirovibrio* (*2, 3*) and *Vampirococcus* (*4*) insert cytoskeletal protrusions, ‘fangs’, which extract the cytoplasm from the attacked cell. Some members of the genus *Cytophaga* are ‘cell eaters’ (*5, 6*), and *Lysobacter* are ‘lysers of bacteria’ (*7*). These and members of the Myxococcales are social organisms which hunt in packs (*8, 9*). Many of these organisms can also subsist as saprotrophs, and thus are facultative predators (*10*), in contrast to *Vampirovibrio* and *Bdellovibrio*, which are obligate predators (*11*). Most of what we know about the physiology, growth, and activity of predatory bacteria has been learned from laboratory studies, because of the difficulty of measuring taxon-specific bacterial activity *in situ*.

Predators are thought to be functionally significant in microbial food webs, but quantitative estimates *in situ* have been very difficult to obtain. It is possible to use fluorescent markers and plate counts to estimate growth rates of predators in artificial media (*12*), but applying such approaches in the field is challenging. For example, it is known that phages prey upon cyanobacteria in rice paddy soils, but the rates of predation are unknown (*13*). Experimental manipulations of soil protozoa in mesocosm studies demonstrate the importance of these eukaryotic predators for nitrogen cycling (*14*) and for decomposition of plant litter (*15*), but the quantitative impacts on these ecosystem processes under field conditions are difficult to measure experimentally. Varying environmental conditions also influence predator-prey interactions: changing moisture content alters soil connectivity, stabilizing or destabilizing predator-prey dynamics (*16*). It is important that predator activity and growth be measured under realistic and varying conditions.

Although protists (*17*), rotifers (*18*), nematodes (*19*), and phages (*11, 20, 21*) are thought to function as the dominant predators in microbiomes, predatory bacteria are common in both soil (*8, 22*) and aquatic (*23*) systems. But beyond their common occurrence in these habitats, we know little of their activity in the wild, how rapidly they grow, their functional significance in food webs, and how they respond to enrichment at the base of the food web through substrate additions.

DNA sequencing and other ‘omics techniques can provide detailed information on the composition and functional potential of the microbiome (*24*), but most measurements of *in situ* bacterial growth rates lack taxonomic resolution and are conducted at the scale of the entire microbial assemblage (*25, 26*). Such aggregate measurements mask the contributions of genetically and functionally distinct populations. Even in macroscopic assemblages, taxa are known to vary in their influences on ecosystem processes (*27*). Techniques that combine isotopes and genetic sequencing hold promise for parsing the contributions of individual microbial taxa to interactions within microbial assemblages and to biogeochemical processes (*28, 29*).

Here, we synthesized measurements using quantitative stable isotope probing (qSIP), a technique that quantifies the isotopic composition of DNA after exposure to an isotope tracer (*30*). qSIP with ^13^C-labeled organic matter tracks the rate of labeled carbon assimilation into DNA, and qSIP using ^18^O-water tracks the incorporation of oxygen from water into DNA. Recovery of the isotope tracer in taxon-specific DNA sequences reflects rates of growth and carbon assimilation of individual microbial taxa (*28, 31*). The survey conducted here included qSIP measurements conducted in natural microbial assemblages from sites in North America, including 14 soils (one arctic, one boreal, 11 temperate, and one tropical), along with one temperate stream (Fig 1, Table 1). We evaluated this dataset to compare rates of growth by predatory and non-predatory bacteria, and their responses to substrate addition.

**Figure 1.**
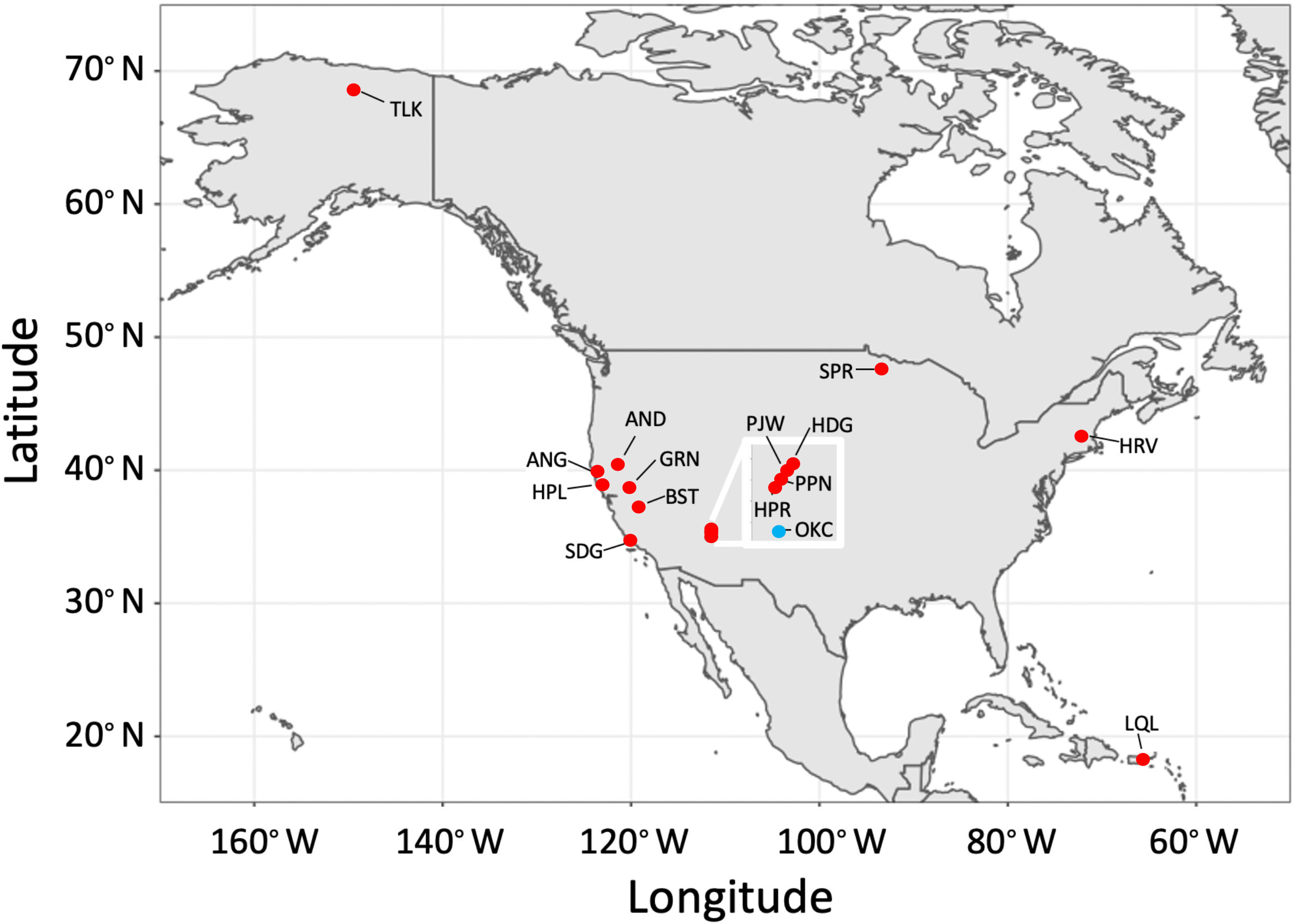
Location of sites included in our meta-analysis of growth rates of predatory and non-predatory bacteria. Additional site information and abbreviations are shown in Table 1. Inset shows a cluster of sites in Arizona (box scale is 1 × 1∘).

**Table 1.**
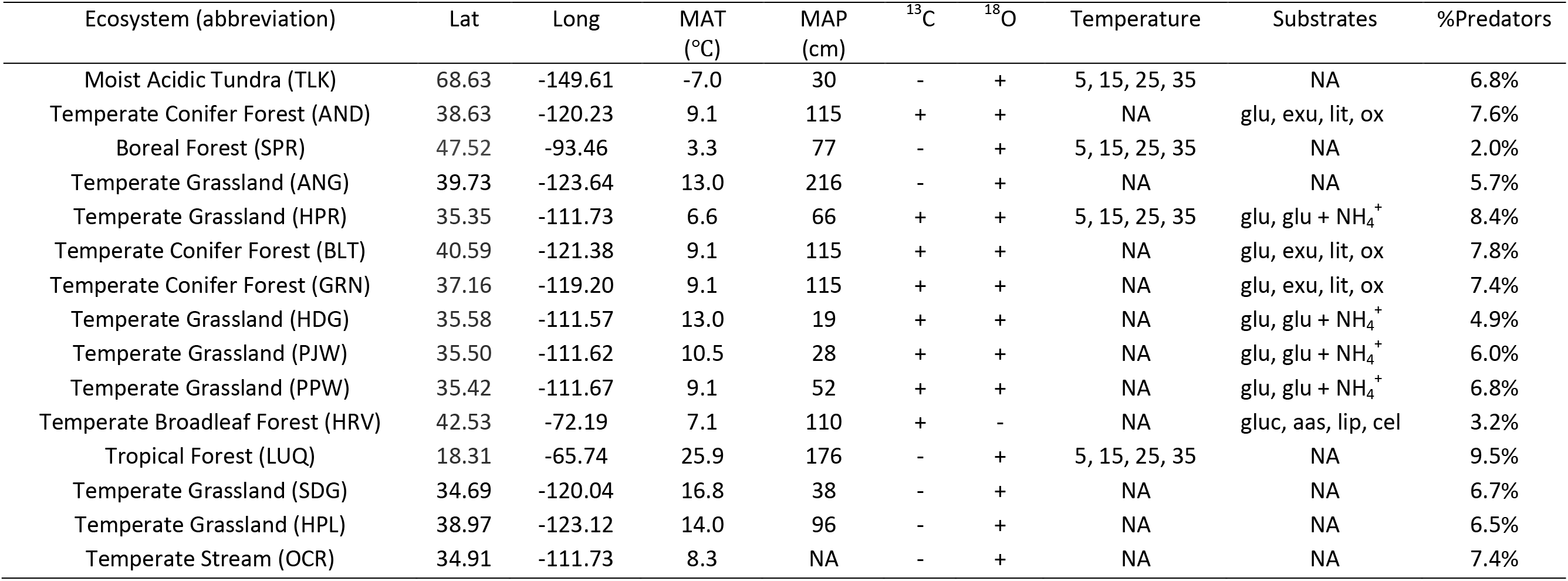
Site description. The columns “Temperature” and “Substrates” indicate experimental treatments applied during the qSIP assay, with temperatures in degrees C and substrates compared to a control with no added substrate. glucose (glu), glucose with ammonium (glu + NH_4_^+^), a mixture of compounds simulating root exudates (exu)(*62*), plant litter, and oxalic acid. Temperature indicates experimental incubation temperatures.

| Ecosystem (abbreviation) | Lat | Long | MAT<br>(°C) | MAP<br>(cm) | <sup>13</sup>C | <sup>18</sup>O | Temperature | Substrates | %Predators |
| --- | --- | --- | --- | --- | --- | --- | --- | --- | --- |
| Moist Acidic Tundra (TLK) | 68.63 | -149.61 | -7.0 | 30 | - | + | 5, 15, 25, 35 | NA | 6.8% |
| Temperate Conifer Forest (AND) | 38.63 | -120.23 | 9.1 | 115 | + | + | NA | glu, exu, lit, ox | 7.6% |
| Boreal Forest (SPR) | 47.52 | -93.46 | 3.3 | 77 | - | + | 5, 15, 25, 35 | NA | 2.0% |
| Temperate Grassland (ANG) | 39.73 | -123.64 | 13.0 | 216 | - | + | NA | NA | 5.7% |
| Temperate Grassland (HPR) | 35.35 | -111.73 | 6.6 | 66 | + | + | 5, 15, 25, 35 | glu, glu + NH<sub>4</sub><sup>+</sup> | 8.4% |
| Temperate Conifer Forest (BLT) | 40.59 | -121.38 | 9.1 | 115 | + | + | NA | glu, exu, lit, ox | 7.8% |
| Temperate Conifer Forest (GRN) | 37.16 | -119.20 | 9.1 | 115 | + | + | NA | glu, exu, lit, ox | 7.4% |
| Temperate Grassland (HDG) | 35.58 | -111.57 | 13.0 | 19 | + | + | NA | glu, glu + NH<sub>4</sub><sup>+</sup> | 4.9% |
| Temperate Grassland (PJW) | 35.50 | -111.62 | 10.5 | 28 | + | + | NA | glu, glu + NH<sub>4</sub><sup>+</sup> | 6.0% |
| Temperate Grassland (PPW) | 35.42 | -111.67 | 9.1 | 52 | + | + | NA | glu, glu + NH<sub>4</sub><sup>+</sup> | 6.8% |
| Temperate Broadleaf Forest (HRV) | 42.53 | -72.19 | 7.1 | 110 | + | - | NA | gluc, aas, lip, cel | 3.2% |
| Tropical Forest (LUQ) | 18.31 | -65.74 | 25.9 | 176 | - | + | 5, 15, 25, 35 | NA | 9.5% |
| Temperate Grassland (SDG) | 34.69 | -120.04 | 16.8 | 38 | - | + | NA | NA | 6.7% |
| Temperate Grassland (HPL) | 38.97 | -123.12 | 14.0 | 96 | - | + | NA | NA | 6.5% |
| Temperate Stream (OCR) | 34.91 | -111.73 | 8.3 | NA | - | + | NA | NA | 7.4% |

## Methods

Atom fraction excess (AFE) values for ^18^O and ^13^C were extracted from qSIP measurements. AFE values were used to estimate bacterial growth rates based on ^18^O assimilation from ^18^O-labeled water, and ^13^C assimilation rate from ^13^C-labeled organic substrates, using methods described in (*30, 32, 33*). All qSIP measurements involved parallel incubations with samples receiving either isotopically labeled (e.g., 97 atom % ^18^O-H_2_O, 99 atom % ^13^C -glucose) or unlabeled substrates (e.g., water with natural abundance ^18^O, or glucose with natural abundance ^13^C). Incubations lasted for 7.1 ± 1.8 days (average ± SD). After each incubation, DNA was extracted and subject to density separation via isopycnic centrifugation. Density fractions were collected, the 16S rRNA gene was sequenced, and the total abundance of 16S rRNA gene copies in each fraction was quantified using qPCR. Quantitative stable isotope probing calculations were then applied to estimate the atom fraction excess ^18^O or ^13^C of each sequenced taxon (*30, 31*). 16S rRNA amplicon sequence data synthesized here have been deposited at NCBI under product IDs PRJNA649787, PRJNA649546, PRJNA649571, PRJNA649802, PRJNA 669516, PRJNA701328, and PRJNA702085.

Across the 15 sites, multiple qSIP measurements were conducted, including experiments within each site. Across all sites and experimental treatments, there were a total of 82 qSIP datasets, and each dataset contained estimates of ^18^O or ^13^C AFE for hundreds of bacterial taxa from a particular site and under a given experimental treatment. The identities of bacterial taxa were used to assign taxa to bacterial groups known to be capable of predation or to non-predatory taxa. Predators were assigned based on belonging to one of six bacterial taxonomic groups known to exhibit predatory behavior: Bdellovibrionales, Cytophagales, *Lysobacter*, Myxococcales, Streptomycetales, and Vampirovibrionales. We recognize that assuming these taxa are unambiguously predatory based on their taxonomic assignment is uncertain. In particular, the facultative groups are known to vary in substrate utilization; the designation of “facultative” acknowledges the range of feeding behaviors exhibited by large groups such as the Cytophagales (*34*), Streptomycetales (*35*), and Myxococcales (*34*). Not all taxa in these groups have been documented to be predatory; we use such broad groups because finer divisions are not available for the trophic behaviors of these organisms. Also, our approach relies on taxonomic assignments based on 16S rRNA gene sequences, which can be unreliable for delineating species or strain (*36*). In 98% of cases, we were able to assign taxa to possible predator groups based on name occurrences in Class, Order, or Family, the higher levels of taxonomic resolution where 16S rRNA gene assignments have been found to be more robust (*37*).

Growth rates were estimated using ^18^O qSIP after accounting for potential differences in the sources of ^18^O among organisms functioning at different trophic levels. qSIP-derived estimates of growth rate using ^18^O-H_2_O begin with the observation that some of the oxygen in DNA is derived from the oxygen in water, so the assimilation of ^18^O from water into DNA reflects its rate of replication, a proxy for cellular growth (*38*). Ribose sugars, nitrogenous bases, and phosphate (*39*) all acquire oxygen from water (*38*). Therefore, the DNA of predators will likely contain oxygen both from water in their growth environment as well as from cellular constituents of prey; these two potential sources of ^18^O in predator DNA may or may not be additive.

To distinguish between these two sources, we compared ^18^O versus ^13^C enrichment in predatory taxa—since many of our SIP studies included treatments with both labeled water and labeled organic C substrates (Table 1). It is standard in food web studies using isotope tracers to treat the ^13^C isotope composition of predator taxa as a conservative indicator of the ^13^C composition of their prey (*40*). The qSIP datasets we evaluated included a subset of dual-isotope measurements, where both ^18^O and ^13^C were determined in parallel experiments with ^18^O-labeled H_2_O and ^13^C-labeled carbon substrates. These measurements occurred in separate incubations, with identical conditions and resource availability, but with different isotope labels applied: in one case ^18^O water was added with a natural abundance carbon substrate, and in the other the carbon substrate was ^13^C-labeled while the added water was at natural abundance ^18^O. With these parallel measurements, we were able to estimate both the ^13^C and ^18^O for multiple taxa.

Across 5 sites and 12 experiments, there were 2197 simultaneous measurements of ^13^C and ^18^O, including 2060 cases of non-predatory taxa and 137 cases of predatory taxa. We evaluated the relationships between ^18^O and ^13^C for both predator and non-predator taxa, reasoning that the two sources of ^18^O to predators (compared to one source for non-predators) would result in predator DNA that was relatively higher in ^18^O compared to ^13^C, to the extent these sources were additive. As expected, for a given value of ^13^C, predator taxa had higher values of ^18^O than non-predator taxa (Figure S1). We used the difference in the relationships (model II linear regressions) between ^18^O vs ^13^C for predators and prey (Figure S1) to predict what the ^18^O composition of predator taxa would have been based on growth on ^18^O-labeled H_2_O alone. This approach resulted in the following correction, which was applied to all predator taxa in the dataset:

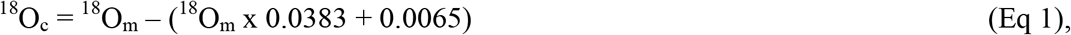

where ^18^O_m_ is the measured predator AFE value and ^18^O_c_ is the adjusted value.

This approach allowed us to avoid overestimating growth rates of predators because of their dual ^18^O sources and helps ensure values of predator and prey AFE ^18^O were comparable. For non-predator taxa, we used the measured qSIP ^18^O AFE value as the estimate of ^18^O assimilation from ^18^O-H_2_O, the standard approach in ^18^O-qSIP studies (*31, 38*). An additional consideration is that oxygen concentration can affect ^18^O assimilation from labeled water (*41*). Although oxygen concentrations were not measured in the incubations, for the Mixed Conifer, Ponderosa, Pinyon-Juniper, and Grassland sites included here, median final CO_2_ concentrations were 0.31% (0.81%, 95^th^ percentile) (*42*), which translates to a small change in atmospheric O_2_, and suggests that oxygen depletion during the incubations was unlikely to have reached levels shown to affect ^18^O assimilation from labeled water (*41*).

Experiments with ^18^O were conducted by adding 97 atom% ^18^O H_2_O to the experimental system and incubating for several days. Because background levels of unlabeled water were present, the ^18^O composition of water in each incubation was determined as a function of the amount of 97 atom % ^18^O water added and the amount of background water. Relative growth rate for each taxon was estimated according to equation 7 from ref (*31*), using AFE ^18^O_h_ of individual bacterial taxa, the AFE ^18^O of water during the incubation, and the duration of the incubation in days.

We compared AFE, growth rates, and carbon assimilation rates of predatory and non-predatory bacteria using meta-analysis (metafor package in R (*43*)), using the log ratios of predator:non-predator as the metric of difference between trophic strategies. This analysis was tested across all sites, treatments, measurement conditions, and tracers. Because some sites included experiments with both ^18^O and ^13^C tracers, isotope treatment was nested within site to preserve independence. For all analyses, site was included as a random effect, because sites included multiple effect sizes which were not independent from each other. Computing multiple estimates with the same control group induces dependency on sampling errors, requiring the use of a variance-covariance matrix in the analysis (*44*). We computed the covariance in log response ratios as

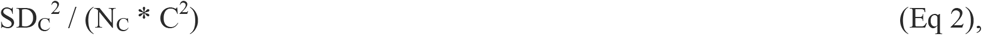

where SD_C_ is the standard deviation of the control group, C is the mean, and N_C_ is the sample size.

We tested for the effect of predator identity on AFE, growth rate, and carbon assimilation rate. Predator identity was evaluated by taxonomic assignment and functional group: obligate predators (Bdellovibrionales and Vampirovibrionales) and facultative predators (Cytophagales, *Lysobacter*, Myxococcales, and Streptomycetales). The effect of predator identity was nested within experiment, because multiple predator groups occurred in the same dataset, so their assimilation rates were not independent of each other.

We used a similar meta-analysis model to evaluate the influence of added carbon substrates on the relationship between growth rates of predatory and non-predatory bacterial taxa. 24 of the compiled qSIP datasets included experimental substrate additions, in which ^18^O-H_2_O qSIP was conducted in soils amended with various carbon substrates compared to a control. Substrates included glucose (6 experiments), oxalic acid (2), ground plant litter (6), a mixture of glucose and ammonium (4), and a mixture of sugars, organic acids, and amino acids simulating root exudates (6). Across all substrate addition experiments and predator taxonomic groups, there were 113 log ratios comparing predator and non-predator growth rates with substrates added, and 187 log ratios comparing predator and non-predator growth rates without substrates added. (The compiled dataset also included experimental manipulations of temperature and of leaf litter species, but the sample sizes were too small to evaluate these as potential drivers.) We evaluated the effect of substrate addition on the growth rates of predators using models with both predator identity and substrate as moderators.

## Results and Discussion

Bacterial taxa identified as potentially predatory were detected at all sites and amounted to 7.4 ± 6.0% of taxa detected at each site (median ± standard deviation). We refer to these as “predatory bacteria” henceforth, acknowledging the limitations of that designation based on 16S rRNA sequence variation — see methods. Most of the predatory bacteria detected were facultative, with 64.7% from the order Myxococcales, 16% from the class Cytophagia, and 9.2% from the order Streptomycetales. 8% were obligate predatory bacteria, with 7.0% from the order Bdellovibrionales, and 1.0% from the order Vampirovibrionales.

Across all sites and experiments, predatory bacteria assimilated isotope tracer into their DNA at rates 23.1 ± 7.0% higher than non-predatory bacteria (meta-analysis, P=0.002, N=407, Figure 2). Climate appeared to have little discernable influence on the differential isotope uptake between predatory and non-predatory bacteria, with weak and non-significant relationships across sites for mean annual temperature (P=0.336) and for precipitation (P=0.738). Soil pH (P=0.871) and soil water content (P=0.165) also had no statistically discernable influence on the relative isotope assimilation between predators and non-predators. Given the current design (15 sites), power may have been limited for detecting such environmental effects.

**Figure 2.**
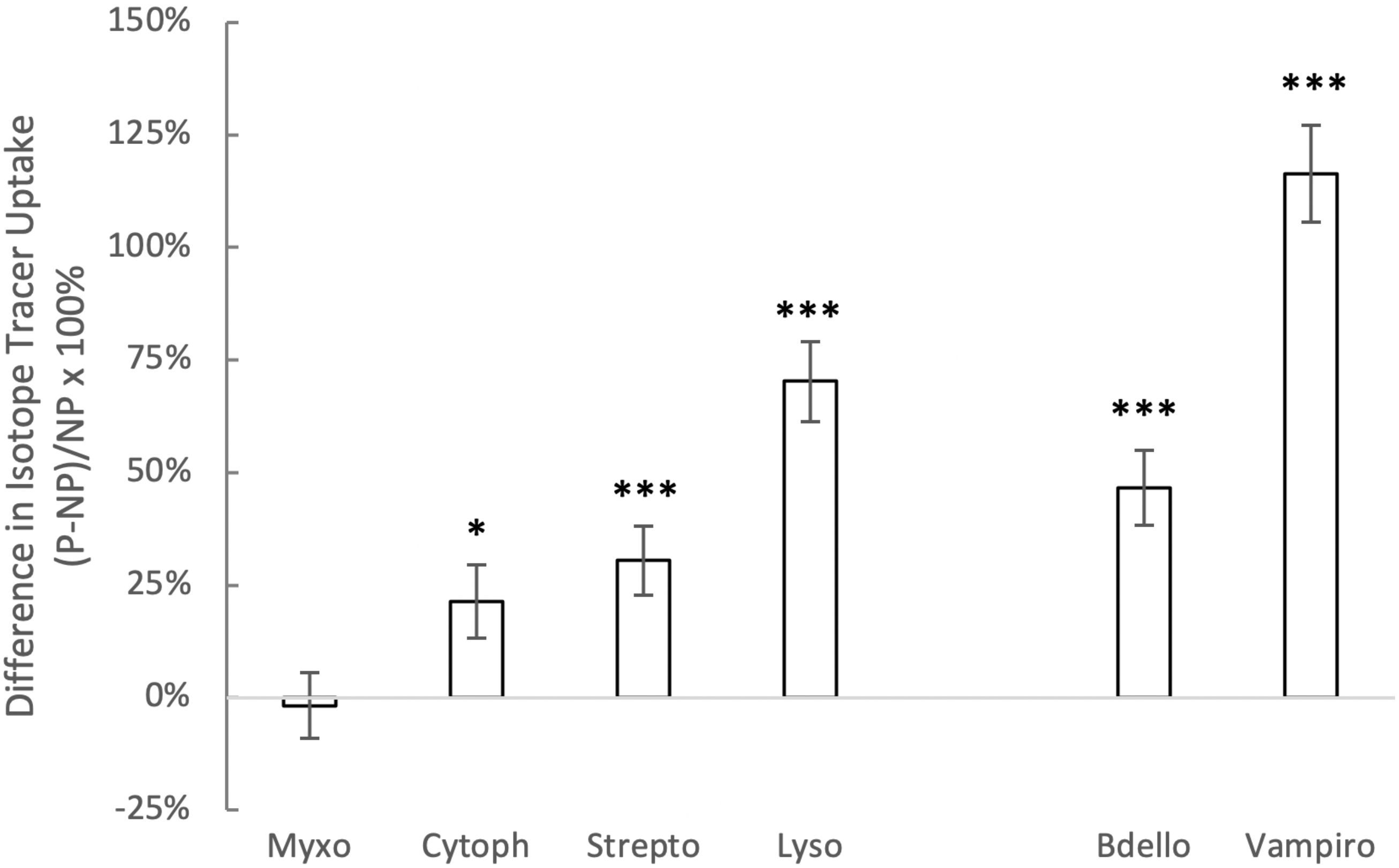
Difference in isotope tracer uptake (^18^O and ^13^C) between predatory and non-predatory bacteria. From left to right, the first four taxa are facultative predators and the last two are obligate predators. Symbols are means ± standard errors of the mean. Predator groups (and numbers of experiments in which they occurred) were Bdellovibrionales (n=71), Cytophagia (n=71), Lysobacter (48), Myxococcales (106), Streptomycetaceae (86), and Vampirovibrionales (25). Asterisks indicate cases where means were significantly higher than zero (* P<0.05 and *** P<0.001).

Predator identity significantly influenced isotope assimilation (P<0.0001, Figure 2): although both obligate and facultative predators assimilated the isotope tracers at rates higher than non-predatory bacteria, the difference was larger for obligate (57.7 ± 8.4%, P<0.001) compared to facultative (17.6 ± 7.1%, P=0.019) predatory bacteria. Finer resolution revealed taxon-specific patterns, with especially high isotope uptake in the members of the obligate predator order Vampirovibrionales (*2, 3*), and in the genus *Lysobacter*, which is known to exhibit wolf-pack type predation (*7–9*). Isotope uptake was also higher in the Bdellovibrionales, Streptomycetaceae, and Cytophagia, whereas rates of isotope uptake for the Myxococcales, many of which are thought to function as saprotrophs (*10*), were similar to rates of non-predators. The higher values of recovery of ^13^C and ^18^O in the DNA of bacterial predators indicates relatively high rates of element flux through bacterial predators in the microbial food webs represented in this 15-site survey.

Across the 15 sites, bacterial growth rates were log-normally distributed, with a median growth rate of 0.035 d^−1^, and 95% confidence from 0.003 to 0.198 d^−1^, a range consistent with past estimates (*31*). The difference in growth rates between predators and non-predators was higher for obligate predators than for facultative predators (Figure 3A). The pattern held for rates of C uptake from ^13^C-labeled substrates: obligate predators had significantly higher C uptake compared to facultative predators and non-predatory bacteria (Figure 3B).

**Figure 3.**
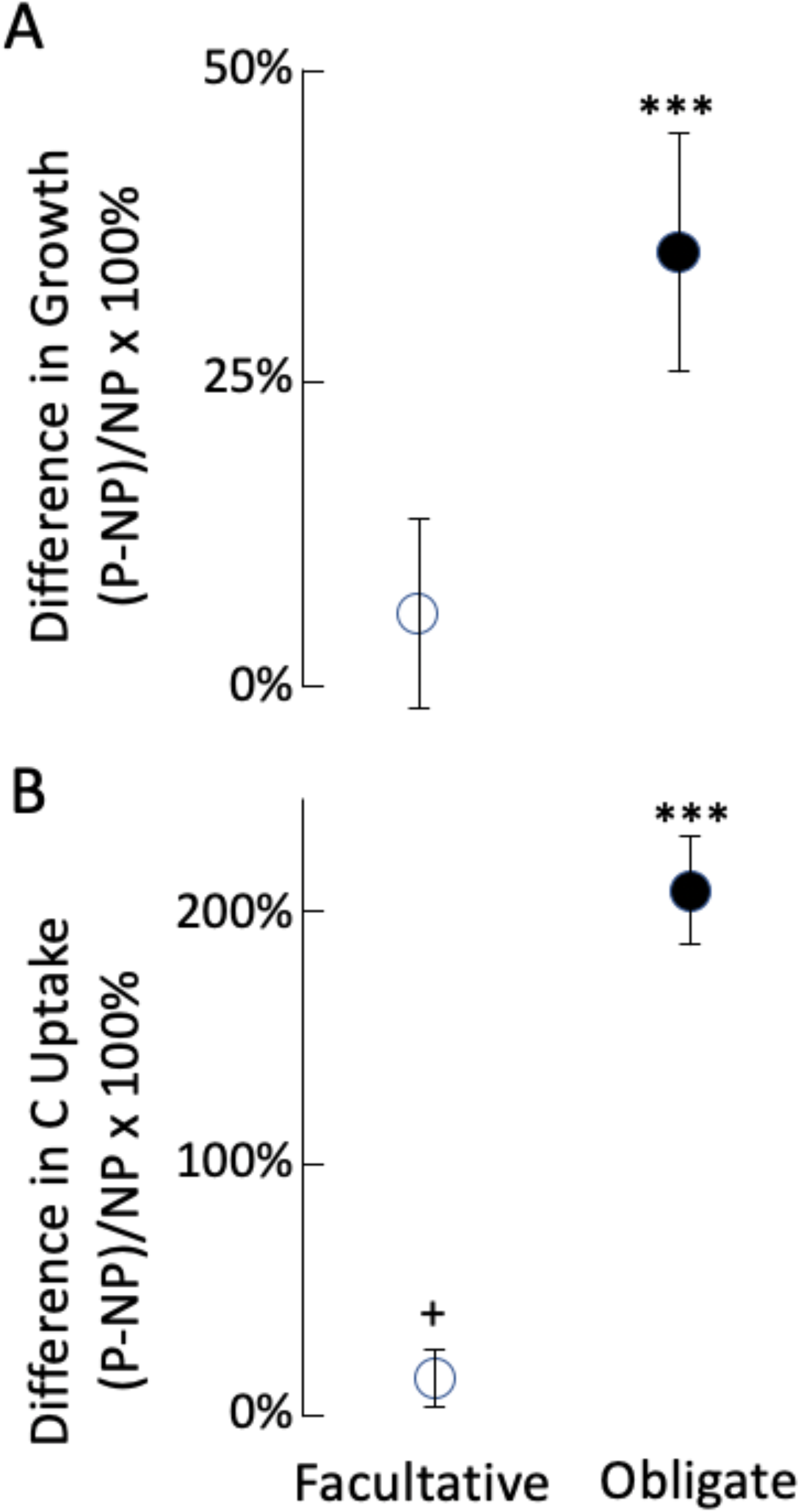
Relative difference in predator growth rate (A) and ^13^C uptake rate (B) compared to non-predators. Values are shown separately for facultative (open symbols) and obligate (filled symbols) predators. Symbols are means ± standard errors of the mean. Statistical results from meta-analysis: *** indicates P<0.001, and + indicates P<0.100.

Adding a source of energy for heterotrophs, in the form of carbon substrates, disproportionately stimulated growth rates of obligate predatory bacteria, whereas responses were indistinguishable between facultative predatory and non-predatory bacteria (Fig. 4). This indicates that higher productivity increases top down (predator-mediated) control in food webs, that added energy disproportionately flows to the predator trophic level, and that predators exhibit functional responses to shifts in prey resource availability. These findings are consistent with long-standing ecological theory that predicts the functional importance of predators increases with productivity (*45–47*), theory that also has support in macroscopic food webs (*48, 49*), and is consistent with observations in polar ocean systems where boom-bust cycles suggest viral response to increased algal productivity (*50*). The similar response of obligate predators from phylogenetically distant clades (i.e., protebacteria Bdellovibrionales and cyanobacteria Vampirovibrionales) implies that the mode of feeding determines response. As such, similar results may be expected for other obligate predatory clades such as the widely distributed marine clade OM27 (Deltaproteobacteria) and family Halobacteriovoraceae. Across all predator taxa, adding nitrogen and carbon together elicited a larger (P<0.001) growth response (38.6 ± 7.5%) compared to adding carbon alone (19.1 ± 10.4%), indicating that carbon-nitrogen stoichiometry of resources affects energy transfer to predatory bacteria (*51*).

**Figure 4.**
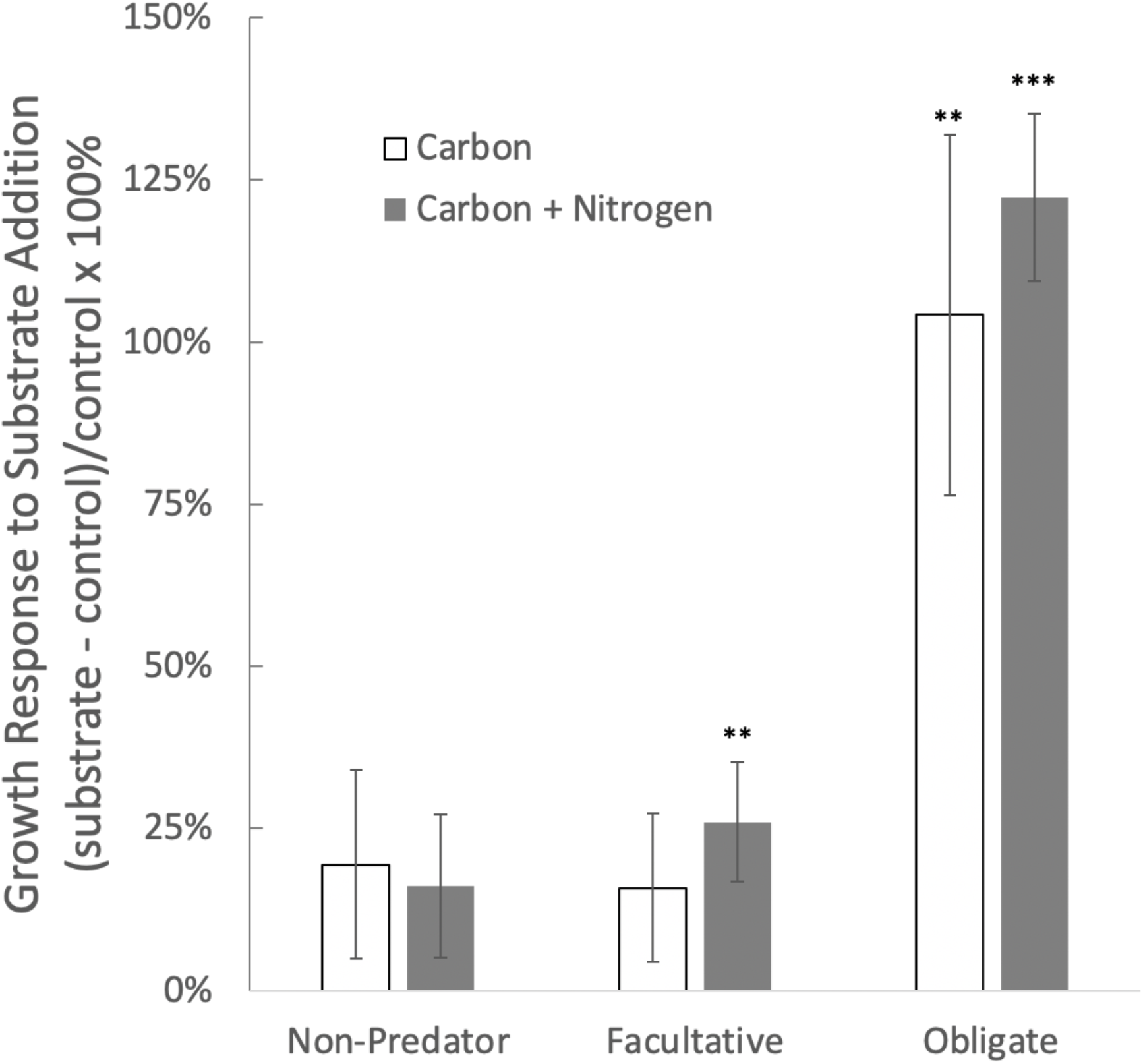
Growth response of predatory and non-predatory bacteria to substrates containing organic carbon or carbon plus nitrogen. Values are means ± SE across 15 sites (Figure 1) where *in situ* growth rates were measured using qSIP with ^18^O-H_2_O. Statistically significant differences from meta-analyses are shown with asterisks, where ** indicates P<0.010 and *** indicates P<0.0001.

Our findings indicate that predatory bacteria are highly active in microbial food webs, synthesizing DNA with elements derived from added isotope tracers at rates higher than non-predatory bacteria, consistent with evidence from experimental microcosms (*52*). These results suggest that bacteria should be considered alongside eukaryotes and viruses as important predators in microbial food webs. Similarly, a recent metagenomic qSIP analysis using a ^13^C-CO_2_ tracer introduced via plant root exudates found that ^13^C recovery in metagenomes associated with putative predator bacteria was comparable to the recovery in viruses and substantially higher than predatory eukaryotes (*53*). Slower growth might be expected if bacterial predators were inactive or dormant, as are many soil microorganisms (*54*). Results presented here indicate that bacterial predators grow, metabolize, and feed at higher rates than most bacteria in the soil food web, and that predatory bacteria may exert top-down effects in microbial food chains. Though our analysis focused on predation, techniques that combine isotopes and gene sequencing can also quantify evidence of other ecological interactions in microbiomes and how they shape carbon flow and nutrient cycling in microbiomes. Multiple signatures of interactions among bacteria have now been identified (*55–57*), informing use of qSIP, metagenomics, and traits to evaluate the functional significance of interactions in diverse microbiomes.

Element flux through the microbiome is central to its functioning, and results from macroecology show how ecological interactions — competition (*58*), mutualism (*59*), and predation (*60, 61*) — strongly influence those fluxes. Evidence presented here synthesizing isotope-enabled microbiome analysis couples predator identity and activity *in situ* and demonstrates that predatory bacteria are highly active in environmental microbiomes, more active than the average bacterial member. Patterns observed across the sites surveyed indicate that top-down trophic interactions are an active force that may structure the composition of element flow in microbiomes and clearly suggests the functional significance of predatory bacteria in microbial food webs.

## Acknowledgements

We appreciate the discussion from participants at the 2020 LLNL ‘Microbes Persist’ Soil Microbiome Scientific Focus Area meeting which inspired this study. This analysis was supported by the U.S. Department of Energy, Office of Biological and Environmental Research, Genomic Science Program (GSP) awards #SCW1632 and DE-SC0020172, and by a Lawrence Fellow award to C.T. through Lawrence Livermore National Laboratory. Studies surveyed in the meta-analysis were funded by DOE GSP awards DE-SC0016207, DE-SC0020172, SCW1024, SCW1590, and by the US National Science Foundation (DEB-1241094, DEB-1645596, DEB-1655357, EAR-1124078). Work at LLNL was performed under the auspices of LLNL under Contract DE-AC52-07NA27344.

## Supplemental Material

Figure S1: Relationship between ^13^C and ^18^O for predator and non-predator taxa from experiments where both ^18^O and ^13^C qSIP were conducted. Lines show major axis model II regression relationships, where models were statistically significant (P<0.001) for both predators: and for non-predators:

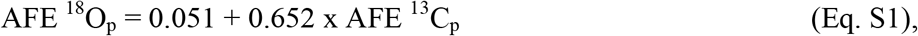

and for non-predators:

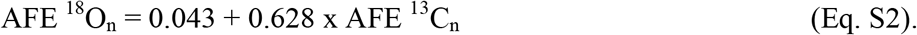

